# Within-Individual Precision Mapping of Brain Networks Exclusively Using Task Data

**DOI:** 10.1101/2025.02.25.640090

**Authors:** Jingnan Du, Maxwell L. Elliott, Joanna Ladopoulou, Mark C. Eldaief, Randy L. Buckner

## Abstract

Precision mapping of brain networks within individuals has become a widely used tool that prevailingly relies on functional connectivity analysis of resting-state data. Here we explored whether networks could be precisely estimated solely using data acquired during active task paradigms. The straightforward strategy involved extracting residualized data after application of a task-based general linear model (GLM) and then applying standard functional connectivity analysis. Functional correlation matrices estimated from task data were highly similar to those derived from traditional resting-state fixation data. The largest factor affecting similarity between correlation matrices was the amount of data. Networks estimated within-individual from task data displayed strong spatial overlap with those estimated from resting-state fixation data and predicted the same triple functional dissociation in independent data. The implications of these findings are that (1) existing task data can be reanalyzed to estimate within-individual network organization, (2) restingstate fixation and task data can be pooled to increase statistical power, and (3) future studies can exclusively acquire task data to both estimate networks and extract task responses. Most broadly, the present results suggest that there is an underlying, stable network architecture that is idiosyncratic to the individual and persists across task states.

---

The organization of human brain networks can be estimated by measuring spontaneous correlations of the blood oxygenation level-dependent (BOLD) signal, a technique referred to as functional connectivity MRI (fcMRI; Biswal et al. 1995; see also Fox and Raichle 2007; Van Dijk et al. 2010; Deco, Jirsa, and McIntosh 2011; Buckner et al. 2013; Murphy et al. 2013; Smith et al. 2013; Power et al. 2014; Lv et al. 2018; Gratton et al. 2020). fcMRI most often uses data acquired during passive states, such as when participants rest with their eyes closed or visually fixate a crosshair. Passive data collection was the procedure adopted early in demonstrations of the technique (Biswal et al. 1995; Greicius et al. 2003) and has remained the mainstay of the field in part because of its simplicity. However, in many instances data have been, or will be, exclusively collected during active task paradigms designed to elicit time locked functional responses, and those data have been considered generally less useful for network analysis.

Some applications require both passive data and active task data. For example, passively acquired data have been extensively employed to estimate brain networks within the idiosyncratic anatomy of individuals (e.g., Laumann et al. 2015; Braga and Buckner, 2017; Gordon et al. 2017; Gordon et al. 2023; Lynch et al. 2024). The networks can then serve as localizers to explore response properties in active task paradigms, including to dissociate functionally distinct side-by-side regions that vary slightly in their positions from one person to the next (e.g., Tobyne et al. 2018; DiNicola, Braga, and Buckner 2020; Du et al. 2024; Edmonds et al. 2024; see also Tavor et al. 2016; Tripathi et al. 2024). Such combination brain mapping experiments often require long (or multiple) sessions given the needed time to separately acquire passive and active task scans. Here we explore the possibility of exclusively using task data to precisely map brain networks within individuals. Serving as the foundation for the present work, prior group-based studies have examined functional connectivity estimates from task acquisitions as compared to passive acquisitions. Fair et al. (2007) specifically raised the possibility of regressing task structure from scans collected during active tasks and then conducting functional connectivity analyses on the residualized data. They observed considerable similarity between the correlation patterns derived from task-regressed data and traditional estimates from passive acquisitions, as well as some differences. Follow-up analyses using multiple acquisition strategies and analysis procedures have found similar results (Cole et al. 2013; Cole et al. 2014; Krienen et al. 2014; Gratton et al. 2016; Xie et al. 2018; Elliott et al. 2019; Ito et al. 2020). Much, but not all, of the region-to-region correlation structure found to be present in resting-state acquisitions is also present in task acquisitions (Krienen et al. 2014; Cole et al. 2014).

However, the portion of the functional correlations sensitive to task states may be sufficient to undermine precision brain mapping (e.g., Geerligs et al. 2015; Salehi et al. 2020; see also Shirer et al. 2012; Gratton et al. 2020). In an influential set of explorations, Salehi and colleagues (2020) illustrated that the borders and exact locations of estimated cortical regions vary as a function of the acquisition task, raising concerns that the estimated regions do not reflect stable, biological subdivisions of cortex. In a critical analysis they demonstrated that task acquisition state could be predicted from the obtained parcellation. Such observations have limited the adoption of task data for the purpose of functional connectivity estimates of brain networks. However, even within the analyses of Salehi et al. there was a high degree of correspondence between parcellation estimates between tasks versus within tasks (see their Figure. 3d). Thus, while there is agreement that the task acquisition state affects the correlation structure of the data, it is an open question of whether the task contribution can be sufficiently minimized, and the commonalities extracted, to estimate stable networks within individuals.

Motivated by the possibility of using existing and future task data for precision brain mapping, we applied and validated a method for estimating brain networks within individuals exclusively using task data. The derived network estimates were remarkably similar to those obtained from traditional resting-state fixation acquisitions. We discovered that, despite historical reservations, task-based data can be effectively used to precisely estimate network organization opening the possibility of novel task designs and increasing efficiency. That is, when analyzed appropriately, task data are sufficient to capture stable estimates of networks within individuals.

## Results

### Functional Correlation Matrices Derived from Task Data Are Highly Similar to Those Derived from Resting-State Acquisitions

Functional connectivity analyses use the region-to-region correlation matrix of the fMRI time series to estimate networks. Our first results examined the similarity between raw correlation matrices derived from task-regressed data (MOT and EPRJ) as compared to traditional resting-state fixation data (FIX). To fully contextualize the correspondence, correlation matrices from independent datasets acquired in the same manner were also examined (e.g., the FIX data were split into independent FIX1 and FIX2 datasets). Four findings were observed (Figure 1).

**Figure 1.**
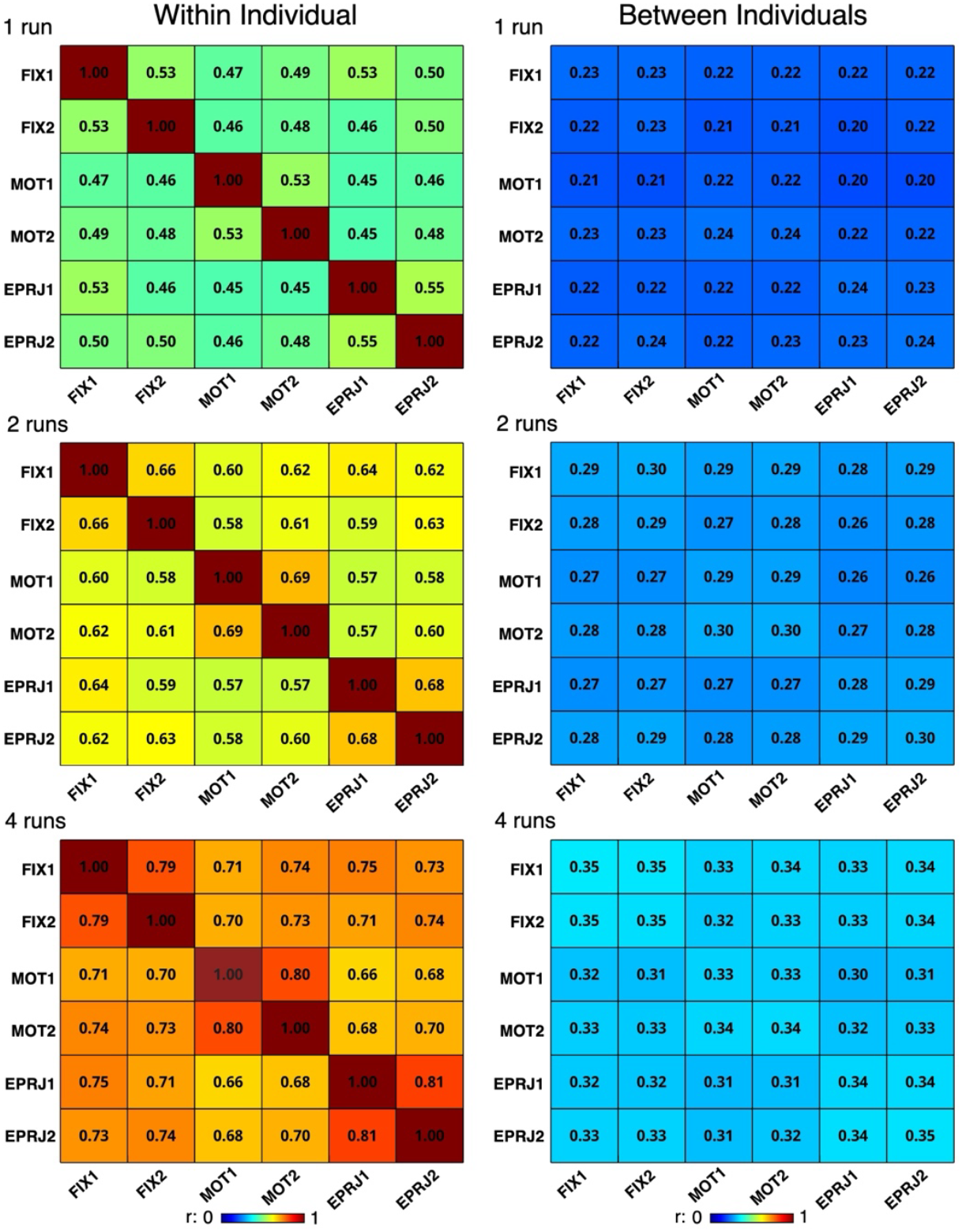
Functional correlation matrices estimated from task-regressed data are similar to those estimated from resting-state fixation data. The similarity estimates between pairs of functional correlation matrices are presented for matched-length data acquired during resting-state fixation (FIX), motor (MOT) and episodic projection (EPRJ) task runs. For each acquisition type, two independent datasets allowed test-retest similarity within an acquisition type (e.g., FIX1 versus FIX2) to be contrasted with similarity between acquisition types (e.g., FIX1 versus EPRJ1). Within-individual similarity was calculated within each participant and then mean averaged across all participants (**Left**). Between-individuals similarity was calculated between each pair of participants and then averaged across all pairs (**Right**). The rows show similarity values for different amounts of data (1, 2, and 4 runs). The colors reflect their correlation strength as noted by the legend below. What is notable is that the correlation values between acquisition types were almost as high as the values within acquisition types for all amounts of data, suggesting that the underlying region-to-region correlation structure is largely, but note entirely, preserved across acquisition conditions. The within-individual similarity values are much higher than the between-individuals similarity values. The primary factor impacting the similarity between functional correlation matrices within individuals was the amount of data.

First, the similarity of the correlation matrices estimated from task-regressed data to the resting-state fixation data was almost as high as any of the data types to themselves. Some variance was related to the task acquisition state, but it was observed to be a relatively small portion of the variance when considering the correlation of same-type data acquisitions to themselves. For example, for the case of 4 runs of data per dataset, the matrices within the same acquisition type correlated with each other with a range of 0.79 to 0.81 (FIX1 versus FIX2, MOT1 versus MOT2 and EPRJ1 versus EPRJ2). The correlations between distinct acquisition types ranged from 0.66 (EPRJ1 versus MOT1) to 0.75 (FIX2 versus EPRJ2). That is, the vast amount of variance in the correlation matrices was similar regardless of the task performed at the time of data acquisition. This result was found whether the task was a low-level sensorimotor task (MOT) or a higher-order task requiring participants to make challenging judgments during extended epochs (EPRJ).

Second, the analyses were repeated for different amounts of data ranging from 1 to 4 runs of data per acquisition type. We found that the similarity between task-regressed and resting-state fixation correlation matrices generalized across all data amounts. The overall correlation strength did change as a function of the amount of data ranging from ∼0.48 when 1 run of data was used (6 min 50 sec) to ∼0.73 when 4 runs of data were used (27 min 20 sec). Thus, as reported previously, increasing the amount of data improves the stability of functional connectivity estimates (Van Dijk et al. 2010; Birn et al. 2013; Laumann et al. 2015; Nee et al. 2019; Lynch et al. 2020).

Third, the similarity of the correlation matrices within the same acquisition type was nearly the same regardless of the acquisition task. For the case of 4 runs of data per dataset, the similarity estimates were 0.53 (FIX1 versus FIX2), 0.53 (MOT1 versus MOT2), and 0.55 (EPRJ1 versus EPRJ2). Thus, no single task acquisition type had an overall benefit over another. This pattern was the same regardless of the amount of data analyzed.

Finally, the similarity of the correlation matrices estimated within individuals was vastly greater than the similarity estimated between individuals. The similarity improved with increasing amounts of data, but the between-individuals correlations for the largest amounts of data (0.30 to 0.35) were still far lower than the within-individual correlations for the smallest amounts of data (0.45 to 0.55). This result suggests that high correlations are not a product of the analysis pipelines and emphasizes that within-individual analyses preserve a great deal of correlation structure that is lost when participants are compared to one another or averaged via anatomical registration (see also Finn et al. 2015; Gratton et al. 2018).

### Functional Correlation Matrices Are Highly Similar Regardless of Smoothing Kernel

To test the effect of spatial smoothing on the similarity of the correlation matrices, we compared smoothing kernel sizes of 0 mm, 2 mm, and 4 mm. This analysis used 4 runs of data (27 min 20 sec) for each acquisition type. Functional correlation matrices were found to be highly stable even without any smoothing (Supplemental Materials). The results also revealed that increasing the smoothing kernel increases the similarity. For example, for the case where there was 0-mm smooth, the matrices correlated within the same acquisition type with a range of 0.74 to 0.75, while for the 4-mm smooth the matrices correlated with a range from 0.83 to 0.85. This suggests that small smoothing kernels, such as 2 mm, are sufficient to capture stable functional connectivity estimates, while potentially preserving spatial specificity in the data. However, there appears to be a consistent (but small) tradeoff between spatial smoothing and reliability.

Relevant to the question of whether the acquisition type has an impact on correlation matrices, the amount of smoothing had little impact on the overall pattern of matrix similarity. The greater the smoothing within the range tested, the higher the similarity between matrices within- and between-task acquisition types, again suggesting that no single acquisition state has a privilege over others.

### Networks Can Be Estimated Robustly Within Individuals Using Task-Regressed Data

The next results explored whether task data are sufficient to generate precision maps of networks that are practical equivalents to traditional maps generated from resting-state fixation data. This is a particularly important question because, while the results above found a high correspondence between the matrices derived from task-regressed data and traditional resting-fixation data, they also revealed clear differences. It is thus an open question whether the similarities between different acquisition conditions are sufficient to drive convergent network estimates or whether network estimates change in substantive ways that might impact experiments exclusively using task data.

Figure 2 provides the answer. When task-regressed data were used exclusively to derive a 15-network Multi-Session Hierarchical Bayesian Model (MS-HBM) estimate, the results were remarkably similar to those generated from traditional resting-state fixation data (e.g., Du et al. 2024). The full parcellation of networks for one representative participant, P2, is displayed in Figure 2. The 15 estimated networks are labeled as follows: Somatomotor-A (SMOT-A), Somatomotor-B (SMOT-B), Premotor-Posterior Parietal Rostral (PM-PPr), Cingulo-Opercular (CG-OP), Salience / Parietal Memory Network (SAL / PMN), Dorsal Attention-A (dATN-A), Dorsal Attention-B (dATN-B), Frontoparietal Network-A (FPN-A), Frontoparietal Network-B (FPN-B), Default Network-A (DN-A), Default Network-B (DN-B), Language (LANG), Visual-Central (VIS-C), Visual-Peripheral (VIS-P), and Auditory (AUD).

**Figure 2.**
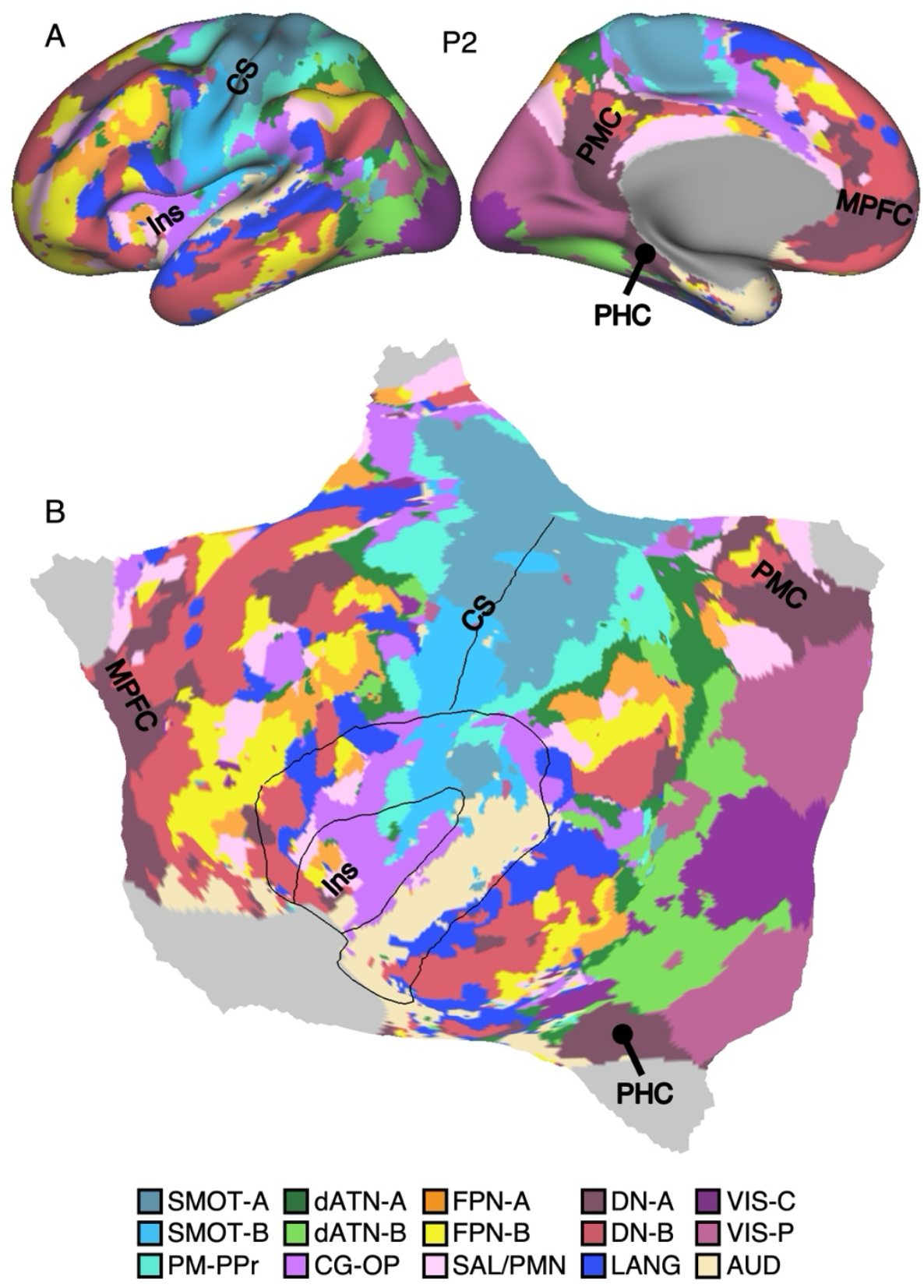
Precision mapping of networks within an individual participant exclusively using task data. Inflated (**A**) and flattened (**B**) surfaces of the left cerebral cortex display the 15-network Multi-Session Hierarchical Bayesian Model (MS-HBM) estimates for participant P2 using task-regressed data. Distinct networks are represented by different colors, with the network labels provided at the bottom. Somatomotor and visual networks possess primarily local organization, whereas other networks possess widely distributed organization across prefrontal, parietal, and temporal association zones. Reference lines illustrate the inner and outer boundaries of the insula (Ins) as well as along the central sulcus (CS). Additional landmarks are posteromedial cortex (PMC), parahippocampal cortex (PHC), and medial PFC (MPFC). The network labels are used similarly throughout the figures. SMOT-A, Somatomotor-A; SMOT-B, Somatomotor-B; PM-PPr, Premotor-Posterior Parietal Rostral; CG-OP, Cingulo-Opercular; SAL, Salience; dATN-A, Dorsal Attention-A; dATN-B, Dorsal Attention-B; FPN-A, Frontoparietal Network-A; FPN-B, Frontoparietal Network-B; DN-A, Default Network-A; DN-B, Default Network-B; LANG, Language; VIS-C, Visual Central; VIS-P, Visual Peripheral; AUD, Auditory.

Figure 3 illustrates a direct comparison of higher-order networks estimated from Fixation Acquisition data versus Task Acquisition data for three representative participants (P2, P6, and P12). This analysis used all available task data (Table 1) and asked the question of whether, independent of the original parcellations estimated in Du et al. (2024) from Fixation Acquisition data, would the same parcellations be obtained in practice if only the task data were used? Precision network maps for all participants are available in the Supplemental Materials. Qualitatively, the network estimates from the two data sources are nearly indistinguishable. The idiosyncratic spatial details varied between individuals but were largely preserved within each individual across datasets under independent acquisition conditions.

**Figure 3.**
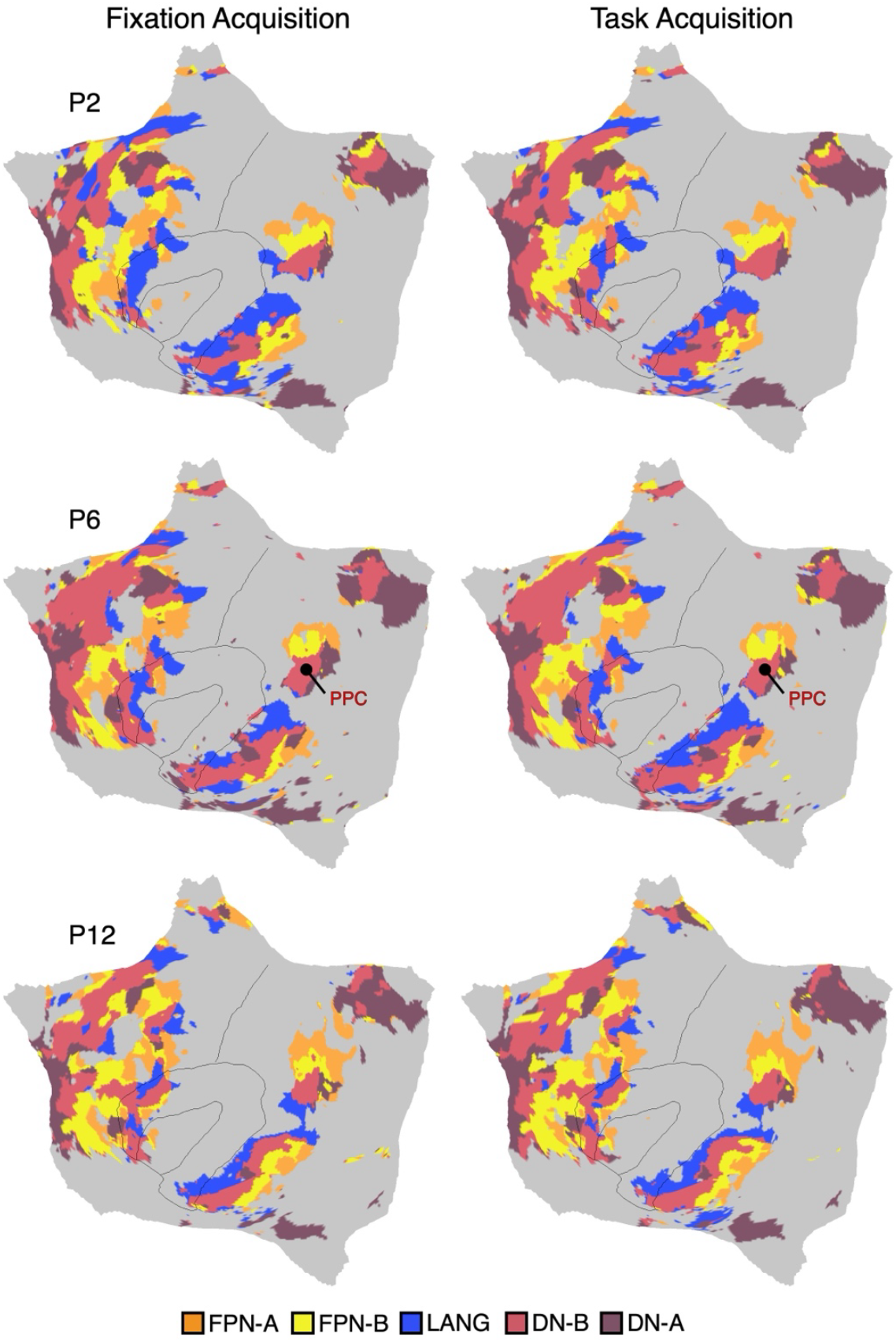
Network estimates are similar between resting-state fixation and task data acquisitions. Higher-order networks from the 15-network MS-HBM estimates are displayed for three representative participants (P2, P6, and P12) for data acquired using traditional resting-state fixation data (**Left, Fixation Acquisition**) versus exclusively using task-regressed data (**Right, Task Acquisition**). The data yielding the network estimate in the left columns is fully independent of the data used in the right columns. The parallel interdigitated distributed networks, FPN-A, FPN-B, LANG, DN-B, and DN-A, are displayed, allowing for direct comparisons of both the broad distributed patterns and idiosyncratic local details. The estimated networks are strikingly similar across both datasets, capturing even idiosyncratic and small regions consistently. A notable example is the juxtaposed spatial segregation between the five networks within the posterior parietal cortex (PPC; labeled for P6). These findings suggest that task-regressed data can effectively estimate brain networks. Similar estimates for all participants are included in the Supplementary Materials. FPN-A, Frontoparietal Network-A; FPN-B, Frontoparietal Network-B; DN-A, Default Network-A; DN-B, Default Network-B; LANG, Language.

**Table 1.** Fixation and task-regressed data used to generate precision maps of networks for each participant.

| ID | FIX<br>(runs) | MOT<br>(runs) | VISME<br>(runs) | ODDBALL<br>(runs) | NBACK<br>(runs) | SENT<br>(runs) | ToM<br>(runs) | EPRJ<br>(runs) | Total<br>Runs |
| --- | --- | --- | --- | --- | --- | --- | --- | --- | --- |
| P2 | 16 | 11 | 5 | 5 | 8 | 6 | 8 | 10 | 53 |
| P3 | 19 | 10 | 5 | 4 | 8 | 6 | 7 | 8 | 48 |
| P4 | 20 | 10 | 5 | 5 | 8 | 5 | 8 | 9 | 50 |
| P6 | 21 | 12 | 5 | 5 | 8 | 6 | 8 | 10 | 54 |
| P7 | 22 | 12 | 5 | 5 | 8 | 6 | 8 | 10 | 54 |
| P8 | 21 | 12 | 5 | 5 | 8 | 6 | 8 | 10 | 54 |
| P9 | 20 | 12 | 5 | 5 | 8 | 6 | 7 | 8 | 51 |
| P12 | 24 | 24 | 5 | 4 | 8 | 11 | 8 | 10 | 70 |
| P13 | 22 | 12 | 5 | 5 | 8 | 6 | 8 | 10 | 54 |
| P14 | 18 | 9 | 5 | 5 | 8 | 6 | 8 | 10 | 51 |
| P15 | 20 | 12 | 5 | 3 | 8 | 6 | 8 | 10 | 52 |

Notes: Numbers represent the number of runs. Participant IDs align with those in Du et al. (2024). Abbreviations: FIX = Resting-state Fixation; MOT = Motor; VISME = Visual Retinotopic Stimulation – Meridians / Eccentricity; ODDBALL = Visual Oddball Detection; NBACK = N-Back Working Memory; SENT = Sentence Processing; ToM = Theory-of-Mind; EPRJ = Episodic Projection.

Quantitative analyses confirmed a high level of correspondence. The overlap percentages of cortical vertices assigned to the same network across the two independent resting-state fixation and task-regressed datasets within the individual was 80.0%: 79.5% (70.5% - 86.7%) for FPN-A, 80.6% (76.2% - 86.5%) for FPN-B, 82.6% (78.4% - 85.9%) for DN-A, 81.4% (75.1% – 87.5%) for DN-B, and 77.1% (67.2% – 84.5%) for LANG. To place these values in context, which on average are ∼80%, we performed a similar analysis for two participants from 30 and 31 sessions of MRI data acquisition under uniform resting-state fixation acquisition. The overlap percentage under such intensive and ideal conditions were 86.4% (81.1% - 87.4% for FPN-A, 82.8% - 90.4% for FPN-B, 88.4% - 88.8% for DN-A, 87.7% – 91.0% for DN-B, and 77.3% – 90.4% for LANG).

Thus, resting-state fixation data and task-regressed data capture a convergent functional architecture of the brain. These findings indicate that task data can be used in practice to estimate precision brain networks that closely align with those estimated from resting-state fixation data.

### Model-Free Seed Region-Based Correlations Confirm Network Estimates

The network estimates in Figures 2 and 3 were derived from a 15-network MS-HBM model. This model assumes a specific number of networks and employs a group prior, which might constrain the network estimates to appear similar between acquisition methods. To explore the network estimates in an unbiased manner and examine how well the model captures underlying within-individual correlation patterns of task-regressed data, we conducted an additional set of analyses using a model-free seed-region based correlation approach. Specifically, we asked whether the correlation patterns from closely juxtaposed seed regions using only the task-regressed data (Task Acquisition) would align with the network borders estimated from the independent resting-state fixation data (Fixation Acquisition). This is a strict test of correspondence because it compares independent data sets each processed with distinct methods that have different assumptions. We focused our analyses on the two adjacent and intertwined networks DN-A and DN-B (Braga and Buckner 2017; DiNicola, Braga and Buckner 2020).

Results were robust and support that the spatial correlation patterns align between the acquisition types (Figure 4). Seed regions placed within the network borders defined from the resting-state fixation data yielded substantially overlapping distributed correlation patterns that align with the independent task-regressed data. In P2, the seed regions were placed in posterior midline zones of the DN-A and DN-B networks (Figure 4 top row). Resulting correlation patterns yielded distinct but closely juxtaposed patterns throughout the cortex that aligned well with the independent estimates of the network borders. In P6, analysis of posterior parietal cortex found similar results (Figure 4 middle row). And in P12, parallel analysis of dorsolateral prefrontal cortex yielded convergent results (Figure 4 bottom row). The Supplementary Materials show results for all participants.

**Figure 4.**
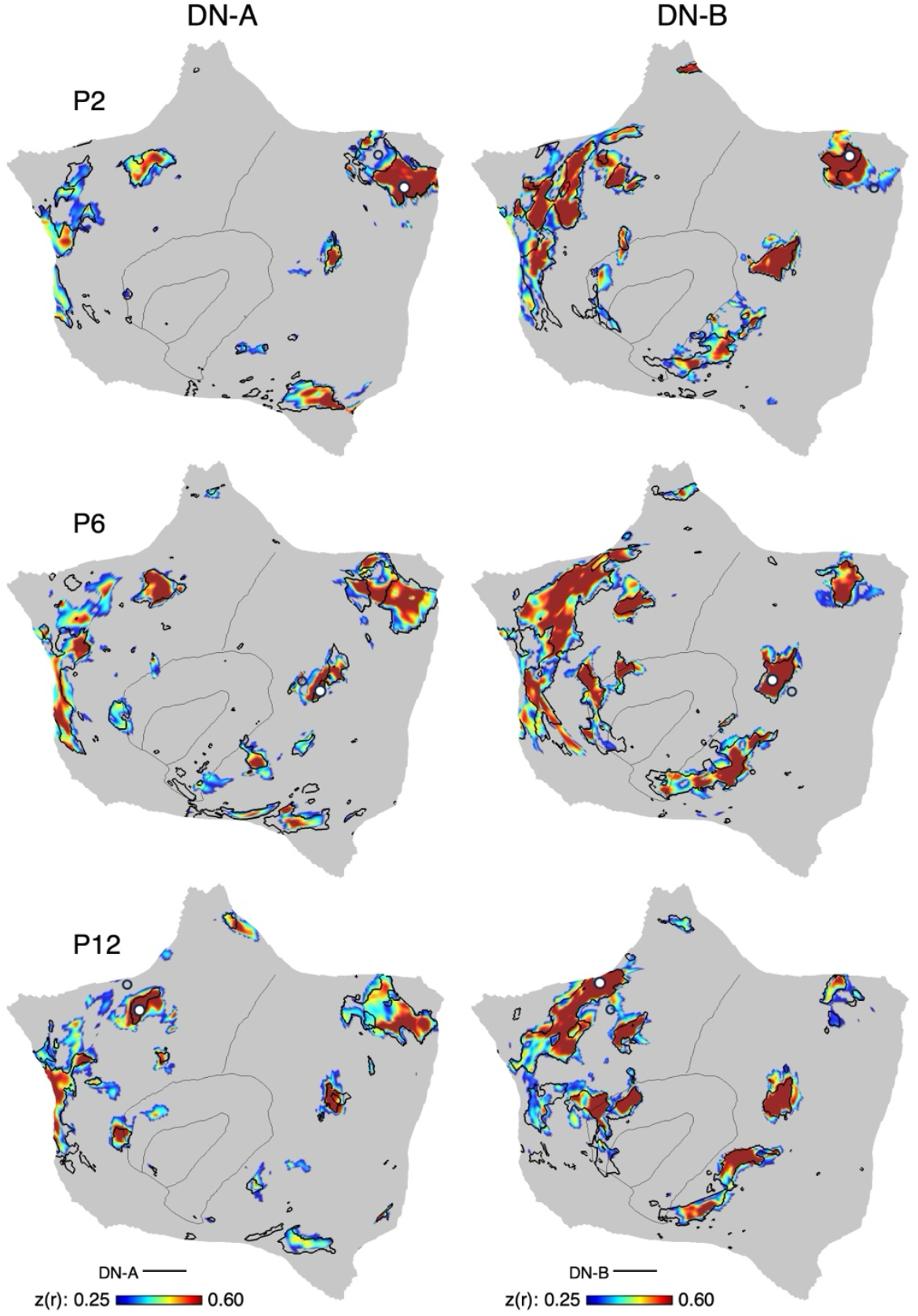
Model-free seed-region correlation maps confirm spatial correspondence between resting-state fixation and task data acquisitions. The correlation maps from individual seed regions placed within networks DN-A and DN-B are displayed for three representative participants (P2, P6, and P12). These correlation maps, based exclusively on task-regressed functional connectivity, are plotted as *z(r)* with the color scale at the bottom. Black outlines show the boundaries of individual-specific networks estimated from the independent resting-state fixation data within the same individual. White-filled circles mark the seed region locations. The positions of the seed regions are moved across association zones between participants to demonstrate that the full distributed extent of DN-A and DN-B can be effectively generated from many component regions. The network borders from resting-state fixation data align well with the spatial correlation properties of the task-regressed data establishing excellent correspondence between resting-state fixation and task-regressed data. Similar estimates for all participants are included in the Supplementary Materials. DN-A, Default Network-A; DN-B, Default Network-B.

### Network Estimates Exclusively from Task-Regressed Data Predict Functional Specificity

Our final result illustrates the practical implications for precision brain mapping under tightly controlled conditions (e.g., matched amounts of Fixation and Task Acquisition data, Table 2). Construct validity was tested by exploring whether task-regressed data yields network estimates that can serve as localizers for prospective explorations. That is, if one collected exclusively task data and estimated networks as a form of within-individual localizer, would the results be similar to traditional approaches? We focused specifically on the functional specialization of three association networks – DN-A, DN-B, and LANG – that are spatially juxtaposed throughout the distributed association zones but established to be functionally distinct.

**Table 2.** Alignment of task and fixation acquisitions for each participant in validity test of functional specificity.

|  | Data Used for Network Estimates |  |  |  |  |  |  | Task Response |  |  |
| --- | --- | --- | --- | --- | --- | --- | --- | --- | --- | --- |
| ID | FIX<br>(min) | TASK<br>(min) | FIX<br>(runs) | MOT<br>(runs) | ODDBALL<br>(runs) | VISME<br>(runs) | NBACK<br>(runs) | SENT<br>(runs) | ToM<br>(runs) | EPRJ<br>(runs) |
| P2 | 109 | 109 | 16 | 11 | 5 | 1 | 0 | 6 | 8 | 10 |
| P3 | 128 | 128 | 19 | 10 | 4 | 5 | 3 | 6 | 7 | 8 |
| P4 | 137 | 138 | 20 | 10 | 5 | 5 | 4 | 5 | 8 | 9 |
| P6 | 143 | 142 | 21 | 12 | 5 | 5 | 2 | 6 | 8 | 10 |
| P7 | 150 | 152 | 22 | 12 | 5 | 5 | 4 | 6 | 8 | 10 |
| P8 | 144 | 143 | 21 | 12 | 5 | 5 | 2 | 6 | 8 | 10 |
| P9 | 137 | 138 | 20 | 12 | 5 | 5 | 1 | 6 | 7 | 8 |
| P12 | 164 | 163 | 24 | 17 | 4 | 5 | 0 | 11 | 8 | 10 |
| P13 | 150 | 152 | 22 | 12 | 5 | 5 | 4 | 6 | 8 | 10 |
| P14 | 123 | 122 | 18 | 9 | 5 | 5 | 2 | 6 | 8 | 10 |
| P15 | 137 | 136 | 20 | 12 | 3 | 5 | 3 | 6 | 8 | 10 |

Notes: Amounts of data from task-regressed (MOT, ODDBALL, VISME and NBACK) runs and resting-state fixation (FIX) runs were matched in length within each individual to define networks. Independent task data (SENT, ToM and EPRJ) were used to examine the task response levels for the independently-defined networks. Numbers indicate the minutes or number of runs for each task, as labelled within the column. Participant IDs align with those in Du et al. (2024). Abbreviations: FIX = Resting-state Fixation; MOT = Motor; ODDBALL = Visual Oddball Detection; VISME = Visual Retinotopic Stimulation – Meridians / Eccentricity; NBACK = N-Back Working Memory; SENT = Sentence Processing; ToM = Theory-of-Mind; EPRJ = Episodic Projection.

Figure 5 displays the functional response levels for these three networks when the networks were defined by resting-state fixation data (Fixation Acquisition) versus exclusively task-regressed data (Task Acquisition). Note that the functional responses plotted are from data that are independent of the data used to define the networks in both cases. DN-A is robustly and preferentially activated for the Episodic Projection (EPRJ) task contrast; DN-B for the Theory-of-Mind (ToM) task contrast; and LANG for the Sentence Processing (SENT) task contrast. Formal statistical analysis using repeated measures ANOVA on network-level task response revealed a significant 3 × 3 interaction between the effects of task contrast and network derived from independent task-regressed data (F(4, 40) = 51.36, *p* < 0.001) and the resting-state fixation data (F(4, 40) = 64.59, *p* < 0.001).

**Figure 5.**
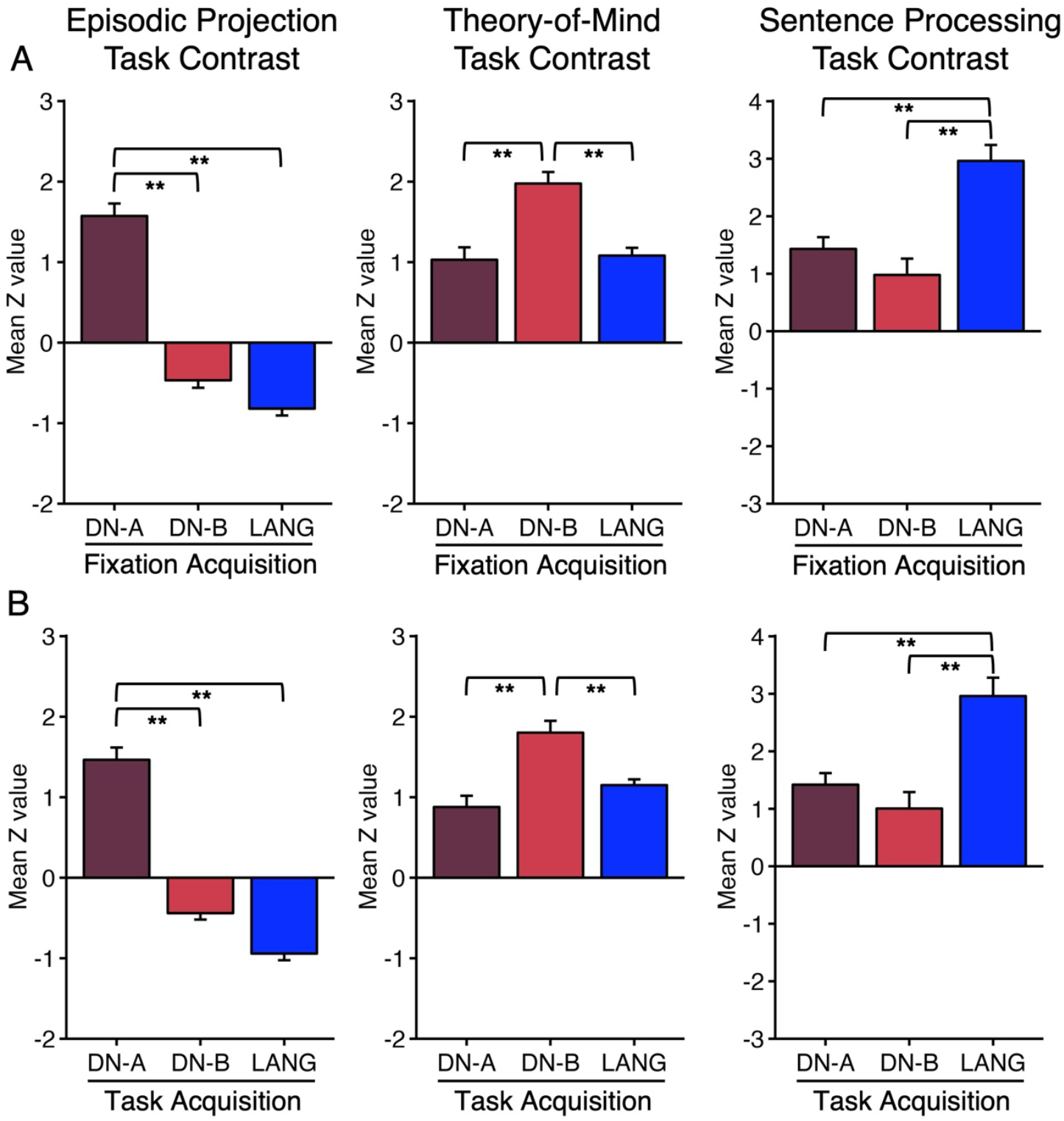
Networks estimated exclusively using task data predict independent task responses as well as traditional resting-state fixation network estimates. As a test of validity, network estimates from traditional resting-state fixation runs (**Top, Fixation Acquisition**) were contrasted with network estimates exclusively using task runs (**Bottom, Task Acquisition**) in their ability to prospectively predict functional dissociations in independent task data. (**Top**) Bar graphs quantify the responses of Episodic Projection (**Left**), Theory-of-Mind (**Middle**), and Sentence Processing task (**Right**) contrasts as mean *z*-values (N = 11) across the multiple *a priori*-defined networks using traditional resting-state fixation data. Note the clear and robust triple dissociation (see Du et al. 2024). (**Bottom**) Similar bar graphs quantify the responses across the multiple *a priori*-defined networks derived exclusively from task-regressed data. The task data used to define the networks was fully independent of the tasks and constructs tested in the bar plots. Each plot displays data from a distinct task contrast; each bar represents a distinct network. The full 3 × 3 interaction (network by task contrast) is significant (*p* < 0.001) for both acquisition types. All pairwise comparisons are also significant for both acquisition types confirming the full triple dissociation. Asterisks indicate a value is significant (** *p* < 0.001).

These results indicate that network estimates from task-regressed data predict functional response properties in independent contrasts similar to parallel analyses using traditional resting-state fixation data.

## Discussion

We provide a procedure for precision mapping brain networks within individuals exclusively using task data. The straightforward approach extends from the observation that region-to-region correlations are similar in task and traditional resting-state fixation data when the contribution of the task structure is minimized (Fair et al. 2007; Cole et al. 2013; Cole et al. 2014; Krienen et al. 2014; Xie et al. 2018; Ito et al. 2020). Building from these prior findings, we regressed out task structure from existing data and then estimated individualized networks, via an MS-HBM model, using the correlation structure in the residualized data. We found that brain networks estimated from task-regressed data are highly similar to estimates from traditional resting-state fixation acquisitions (Figures 2-4) and predict independent functional dissociations (Figure 5). Task data are routinely acquired in research and translational efforts, and future efforts will confront tradeoffs between acquiring data to map network organization and the goal of extracting task responses. The present findings suggest that task data are sufficient to generate precision maps within individuals that can be used for many purposes.

### Within-Individual Precision Mapping of Brain Networks Exclusively Using Task Data

Functional neuroimaging with MRI has commonly used two broad classes of methods – methods that employ task-based contrasts to characterize regions responding to structured task demands and methods that analyze spontaneous fluctuations to detect the intrinsic architecture of brain networks (see Finn 2021 for an historical perspective). Both methods have contributed to important discoveries. However, there has been a practical tension between the two types of methods because they are acquired across different scans. Given limitations on cost and participant burden, imaging sessions typically prioritize one type of data acquisition or strike an uneasy balance. Past data acquired for the purpose of measuring task responses has rarely been used for functional connectivity-based network estimation (for early, uncommon exceptions see Greicius et al. 2003; Greicius et al. 2004; Fair et al. 2007; Andrews-Hanna et al. 2007).

For example, in the Human Connectome Project (HCP), data were acquired over two days with 4 runs of data dedicated to resting-state fixation scans amenable to analyses using functional connectivity and 7 runs dedicated to task paradigms (Van Essen et al. 2013). Similarly, the recent UK Biobank effort, which was substantially constrained by scan length limits, included one run of resting-state fixation data and one run of a face emotion processing task data (Miller et al. 2016). Broadly across the field, including in our laboratory’s efforts, scan runs acquired for the goal of estimating network organization are separated from runs acquired to measure task response.

Building from the insight that the correlation of intrinsic fluctuations between regions is similar for resting-state fixation acquisitions and task acquisitions (after regression of the task structure; Fair et al. 2007), we developed and tested a procedure to estimate brain networks within individuals exclusively from task data. The results were clear. Network estimates from task data converged with traditional estimates from resting-state fixation data. The estimates were qualitatively similar to the eye (Figure 3), quantitatively similar in terms of estimates of overlap, and predicted preferential responses in external task data (Figure 5). That is, the task-regressed network estimates were highly effective functional localizers.

### Implications for Task Designs and Analysis

The present findings have multiple implications. First, large amounts of task data have been collected through local lab efforts and large-scale consortia (e.g., Van Essen et al. 2013; Miller et al. 2016; Gordon et al. 2017; Volkow et al. 2018; Allen et al. 2022; Du et al. 2024). These previously acquired data can be reanalyzed with the present approaches to estimate networks using functional connectivity. Second, for already collected datasets that have both task and resting-state acquisitions, power can be increased for network estimation by combining and using all of the data. Our results (and those across the field broadly) suggest that the amount of data utilized for network estimation is critical. Thus, utilization of all data may allow for sufficient data pooling to stabilize within-individual precision estimates when passively acquired data are limited. For illustration, the supplementary materials show examples where network estimates are stabilized by augmenting the resting-state fixation runs by adding the task-regressed data (Supplementary Materials).

As a counterintuitive possibility, in situations where task and resting-state acquisitions are both acquired, the task data, via the methods proposed here, can be used to estimate the topography of networks within individuals, and then the localized networks used to compute within-and between network correlations in the separate resting-state data. This seemingly odd task design, which to our knowledge has never been conducted, would allow independent data to be used to estimate the location of target regions prior to analysis of region-to-region correlations that may fluctuate over time (e.g., Lynch et al. 2024) or in response to an intervention (Siegel et al. 2024). Using the task-regressed data will allow the networks to be estimated separately from the data used to measure and track changes in the spontaneous correlations between regions in a spatially unbiased manner.

Finally, future data collections might consider to exclusively acquire task data and use that task data to both estimate networks and also to quantify the task response from regions within those networks. This key opportunity is worth elaborating. Many recent findings have demonstrated the importance of taking into consideration the idiosyncratic anatomy of the individual in functional analyses (e.g., Fedorenko et al. 2012; Laumann et al. 2015; Braga and Buckner 2017; Gordon et al. 2017; Smith et al. 2021; Somers et al. 2021; Noyce et al. 2022; Gordon et al. 2023; DiNicola, Sun and Buckner 2023; for further discussion see Gratton and Braga 2021; Laumann, Zorumski, and Dosenbach 2023). In some studies, one set of task runs is used to localize regions and networks, which are then examined in independent task data to quantify response properties within the localized anatomy of the individual. In other studies, network estimates or regional (areal) parcellations are constructed from resting-state fixation or similar data. These estimates are then used as localizers to quantify response properties in independently collected task runs.

The present results suggest that task data can solely be acquired with the task response estimated via the GLM while the task-regressed data are used to estimate networks that act as localizers to focus the task response explorations. For example, if a working memory or language paradigm with multiple conditions was collected, the task-regressed component of the data could be used to localize regions, and then the direct effects of the task paradigms used to explore modulation of those regions. This novel task design is particularly efficient because all of the runs acquired to estimate the task responses can simultaneously be used to estimate the networks (using task residuals) that would normally be discarded in a standard analysis.

## Methods

### Overview

The goal of the analyses was to explore if precision mapping of brain networks can use task data to yield equivalent estimates to those traditionally generated from resting-state fixation data. The analyses proceeded in three phases. First, equivalent amounts of task and resting-state data were used to define correlation matrices that were compared to one another to determine their similarity. The correlation matrices are the basis of functional connectivity analyses so directly contrasting the matrices provides information about correspondence without any assumptions about network models or region definitions. Second, we asked the question of whether task data is sufficient to generate precision maps of networks that are equivalent to traditional maps generated from resting-state fixation data. Correlation matrices from task data were first used to generate within-individual precision maps of networks. The network estimates from the task data were then directly compared to network estimates that used only traditional resting-state fixation data. As a final analysis, construct validity was tested by asking the question of whether task data yield network estimates that could serve as localizers for prospective explorations. Equivalent amounts of data from task acquisitions and resting-state acquisitions were used to define networks. Then, the network estimates were used prospectively to measure functional responses in independent task data that was not used to generate the networks. The critical test was whether known, differential response patterns could be obtained similarly for task-based network estimates as compared to estimates from traditional resting-state acquisitions.

### Participants

Fifteen right-handed native English speakers (labeled P1 to P15) participated for monetary compensation (aged 18 to 34; mean = 22.1 yr, SD = 3.9 yr; 9 female). The data were previously used for traditional resting-state functional connectivity analyses and reported in papers focused on the cerebral cortex (Du et al. 2024), the cerebellum (Saadon-Grosman et al. 2024), the striatum (Kosakowski et al. 2024), and the hippocampus (Angeli et al. 2025). Here the task-based data were reanalyzed with a focus on estimating precision brain networks using task-regressed functional connectivity. Participants with a history of neurological or psychiatric illness were excluded. All participants provided informed consent using protocols approved by the Harvard University Institutional Review Board. Participants were scanned across 8-11 MRI sessions. Each session included multiple resting-state fixation and task-based runs. Four of the fifteen participants were excluded from this study due to insufficient task data, leading to the inclusion of eleven participants (see Table 1; also refer to Table 1 in Du et al. (2024)). Usable resting-state runs ranged from 16 (P2) to 24 (P12) runs. Usable task-based runs ranged from 48 (P3) to 70 (P12) runs (see Table 1).

### MRI Data Acquisition

Details of the methods have previously been reported in Du et al. (2024) with relevant portions repeated here. Scanning was performed at the Harvard Center for Brain Science using a 3-T Prisma^fit^ MRI scanner and a 32-channel phased-array head-neck coil (Siemens Healthineers, Erlangen, Germany). Foam and inflatable padding minimized head movement. Participants viewed a rear-projected display positioned to optimize comfortable viewing. Eyes were video recorded using an Eyelink 1000 Core Plus with Long-Range Mount (SR Research, Ottawa, Ontario, Canada). MRI data quality was monitored during the scan using Framewise Integrated Real-time MRI Monitoring (FIRMM; Dosenbach et al. 2017).

Each participant was scanned across 8-11 sessions most often over 6 to 10 wks. A few participants had longer gaps between the first and last MRI sessions up to one year. Each session involved multiple fMRI runs to be used for functional connectivity analysis, for a total of 16 to 24 resting-state fixation runs, and 48 to 70 task-based runs obtained for each individual. Blood oxygenation level-dependent (BOLD) acquisition parameters were: voxel size = 2.4 mm, TR = 1,000 ms, TE = 33.0 ms, flip-angle = 64°, matrix 92 × 92 × 65 (FOV = 221 × 221), 65 slices covering the full cerebrum and cerebellum. Each resting-state fixation run lasted 7 min 2 sec (422 frames with the first 12 frames removed for T1 equilibration). Dual-gradient-echo B0 fieldmaps were acquired.

High-resolution T1w and T2w structural images were acquired based on the Human Connectome Project (HCP) sequences (Harms et al. 2018). T1w magnetization-prepared rapid gradient echo (MPRAGE) parameters: voxel size = 0.8 mm, TR = 2,500 ms, TE = 1.81, 3.60, 5.39, and 7.18 ms, TI = 1,000 ms, flip-angle = 8°, matrix 320 × 320 × 208, 208 slices, in-plane generalized auto-calibrating partial parallel acquisition (GRAPPA) acceleration = 2. T2w sampling perfection with application-optimized contrasts using different flip angle evolution sequence (SPACE) parameters: voxel size = 0.8 mm, TR=3,200 ms, TE=564 ms, 208 slices, matrix=320 × 320 × 208, 208 slices, in-plane GRAPPA acceleration = 2. Backup, rapid T1w structural images were also obtained using a multi-echo MPRAGE sequence (van der Kouwe et al. 2008): voxel size = 1.2 mm, TR = 2,200 ms, TE = 1.57, 3.39, 5.21, 7.03 ms, TI = 1,100 ms, flip-angle = 7°, matrix 192 × 192 × 176, in-plane GRAPPA acceleration = 4.

*Task-based fMRI Data*. Extensive task-based fMRI data were collected to explore functional response properties (Du et al. 2024). The identical sequence was used for both the task runs and the resting-state fixation runs, ensuring the spatial alignment of the estimated networks from both types of acquisition within each individual. In this study, the task runs were reanalyzed to estimate precision brain networks using standard functional connectivity MRI (fcMRI) network analyses.

### Task Paradigms

#### Tasks Used for Estimating Matrices

In the original study design, fMRI data were acquired during a resting-state fixation task (FIX) to be used for network estimation; independent data were acquired during active task conditions to probe functional response properties (Du et al. 2024). Here the task-based data were reanalyzed to explore whether task-based data can robustly and precisely estimate brain networks using standard within-individual fcMRI network analysis. Detailed descriptions of the tasks including exclusions are provided in Du et al. (2024). Tasks reanalyzed here for functional connectivity analysis included a motor (MOT) task designed to estimate somatomotor topography; an Episodic Projection (EPRJ) task that encouraged participants to remember and imagine the future; a visual oddball detection (ODDBALL) task where participants detected salient, uncommon targets among distractors; a working memory (NBACK) task designed to study cognitive control under memory load; and a visual retinotopic stimulation - meridians / eccentricity task (VISME) designed to map retinotopy.

#### Tasks Used for Test-Retest Reliability and Between-Task Similarity Estimates

The available data allowed for three of the acquisition types to be further divided within each participant into independent test and retest samples yielding six distinct datasets (FIX1, FIX2, MOT1, MOT2, EPRJ1, and EPRJ2), all with equivalent amounts of data (Table 2). To ensure comparability, the MOT and EPRJ runs were trimmed to 410 volumes (6 min 50 sec), exactly matching the amount of data in FIX runs. These six matched datasets enabled contrasts between acquisition types (FIX1 versus MOT2) to be compared to test-retest estimates within the same acquisition type (FIX1 versus FIX2, MOT1 versus MOT2). In this manner, the similarity between matrices of data acquired within the same tasks could serve as a reference for contrasts between matrices of different acquisition types. In addition, having two independent estimates of matrices for each acquisition type allowed us to explore whether any acquisition types generally yielded more reliable estimates. For example, it could be the case that resting-state fixation acquisitions generally yield more similar matrices than matrices derived from the task-based acquisitions.

#### Tasks Used for the Construct Validity Test of Functional Specificity

Three critical tasks were further used to measure functional response within independently defined networks and regions (Sentence Processing, SENT; Theory-of-Mind, ToM; Episodic Projection, EPRJ). Given their importance, these three tasks are described in more detail below. Comprehensive task descriptions, contrast details, and code for all tasks are openly available (https://dataverse.harvard.edu/dataset.xhtml?persistentId=doi:10.7910/DVN/AVB4BW). The Sentence Processing (SENT) task examined domain-specialized processes related to word and phrase-level meaning (Fedorenko et al. 2010; 2012). Participants passively read real sentences or pronounceable nonword strings. After each word or nonword string, a cue appeared for 0.50 sec, prompting participants to press a button with their right index finger. Word and nonword strings were presented in blocks of three strings. Extended fixation blocks appeared at the start of each run and after every fourth string block. The comparison of interest was the contrast between sentence blocks and non-word blocks to activate the LANG network.

The Theory-of-Mind (ToM) tasks investigated domain-specialized processes associated with understanding other people’s mental states (Saxe and Kanwisher 2003; Dodell-Feder et al. 2011; Bruneau et al. 2012; Jacoby et al. 2016). In the False Belief paradigm, participants read brief stories describing a protagonist with a false belief (False Belief condition) or a physical scene (False Photo condition). In the control False Photo condition, participants indicated whether the story was true or false. In the Pain paradigm, stories described emotionally painful situations (Emo Pain condition) and were contrasted with control stories involving physical pain (Phys Pain condition). At the end of each story, participants rated the level of emotional / physical pain experienced by the protagonist during the question period. Stimuli never repeated. The comparison of interest combined the False Belief versus False Photo and Emo Pain versus Phys Pain contrasts extending from DiNicola, Braga, and Buckner (2020).

The Episodic Projection (EPRJ) task examined domain-specialized processes related to remembering the past and imagining the future (prospection) (Andrews-Hanna et al. 2010; DiNicola, Braga, and Buckner 2020). In the target task conditions, the participants read scenarios that oriented them to either a past (Past Self) or a future (Future Self) situation, and simultaneously answered questions about the scenarios. The similarly structured control condition asked the participants about a present situation (Present Self). All scenarios were unique. The comparison of interest combined the Past Self versus Present Self and Future Self versus Present Self contrasts extending from DiNicola, Braga, and Buckner (2020).

## Data Processing

The data were processed using the openly available iProc processing pipeline that preserves spatial details by minimizing blurring and interpolations (described in detail in Braga et al. 2019 and Du et al. 2024; relevant methods are repeated here). For resting-state fixation data, the processed data were taken directly from Du et al. (2024). For task-based data, the data were reprocessed in this study using the same processing steps as for resting-state fixation data.

Data were interpolated to a 1-mm isotropic T1w native-space atlas (with all processing steps composed into a single interpolation) that was then projected using FreeSurfer v6.0.0 to the fsaverage6 cortical surface (40,962 vertices per hemisphere; Fischl et al. 1999). Four transformation matrices were calculated: 1) a motion correction matrix for each volume to the run’s middle volume [linear registration, 6 degrees of freedom (DOF); MCFLIRT, FSL], 2) a matrix for field-map-unwarping the run’s middle volume, correcting for field inhomogeneities caused by susceptibility gradients (FUGUE, FSL), 3) a matrix for registering the field-map-unwarped middle BOLD volume to the within-individual mean BOLD template (12 DOF; FLIRT, FSL), and 4) a matrix for registering the mean BOLD template to the participant’s T1w native-space image which was resampled to 1.0 mm isotropic resolution (6 DOF; using boundary-based registration, FreeSurfer). The individual-specific mean BOLD template was created by averaging all field-map-unwarped middle volumes after being registered to an upsampled 1.2 mm and unwarped mid-volume template (an interim target, selected from a low motion run, typically acquired close to a field map).

For resting-state fixation runs and task-based runs used for functional connectivity analysis, confounding variables including six head motion parameters, whole-brain, ventricular and deep cerebral white matter signals, and their temporal derivatives were calculated from the BOLD data in T1w native space. The signals were regressed out from the BOLD data with 3dTproject, AFNI (Cox et al. 1996; 2012). The residual BOLD data were then bandpass filtered at 0.01–0.1-Hz using 3dBandpass, AFNI (Cox et al. 1996; 2012). For task-based runs used for standard task contrast analysis, only whole-brain signal was regressed out (see DiNicola, Braga and Buckner 2020). The data were then resampled from T1w native-space atlas to the fsaverage6 standardized cortical surface mesh using trilinear interpolation (featuring 40,962 vertices per hemisphere; Fischl et al. 1999) and then surface-smoothed using a 2-mm full-width-at-half-maximum (FWHM) Gaussian kernel for primary analysis and 0-mm and 4-mm FWHM Gaussian kernel for exploration on the impact of spatial smoothing.

### Estimation of Functional Connectivity Matrices

Functional connectivity was estimated by calculating Pearson correlations between time series from pairs of brain regions, using Fisher’s *z*-transformed values for computations, which were then converted back to *r* values for reporting. For resting-state fixation data, this process required no additional steps after processing to calculate the correlations. For task-based data, task structure was removed to allow functional connectivity to be performed on the intrinsic fluctuations unrelated to task events (e.g., trials and blocks). Extending from Fair et al. (2007), we employed regression to remove the task structure from the fMRI data (see also Cole et al. 2013; 2014). Specifically, we used residualized data following standard task-based general linear model (GLM) analysis as applied in Du et al. (2024) and input the resulting task-regressed data (residuals) into the standard fcMRI network analysis pipelines. The benefit of this approach is that it is straightforward and allows use of residualized data that is routinely generated from standard task-based processing pipelines without any additional steps.

### Estimation of Similarity Between Matrices

The broad question of this paper is whether functional connectivity can utilize task data to yield results and network estimates that are equivalent in practice to the traditional use of resting-state fixation data. The first set of analyses examined the equivalence of the correlation matrices directly to determine how similar or different they are when derived from data acquired under different task conditions and following multiple levels of spatial smoothing.

To quantify matrix similarity, we constructed a 1,175 × 1,175 functional connectivity matrix based on 1,175 regions-of-interest (ROIs) uniformly distributed across the two hemispheres (Yeo et al. 2011; Kong et al. 2019). The linearized upper triangles of these symmetrical matrices were used as inputs for calculating similarity values across matrices derived from distinct data sets and following distinct processing choices.

For each task, functional connectivity matrices were calculated for each run and mean averaged to produce a stable functional network matrix, which was subsequently used to calculate the similarity matrix between independent data sets from the same acquisition types and between acquisition types (e.g., FIX1 versus FIX2, FIX1 versus MOT2, MOT1 versus MOT2). The similarity value was directly examined by calculating Pearson correlation among the linearized upper triangles of functional connectivity matrices, resulting in a second order “similarity matrix” (see also Gratton et al. 2018).

As a control and to illustrate the importance of conducting analyses within the idiosyncratic anatomy of the individual participant, we also compared the matrices between individuals rather than within individuals. As will be demonstrated, between-individuals correlations were markedly attenuated reinforcing the importance of conducting analyses within the anatomy of the individual.

### Within-Individual Precision Network Estimates of the Cerebral Cortex

We employed the 15-network MS-HBM previously used by Du et al. (2024) and DiNicola, Sun and Buckner (2023) to estimate networks (derived from the approach developed by Kong et al. 2019). This MS-HBM estimation was applied in small groups, mirroring our previous procedure (see Du et al. 2024 for details). 4 out of 15 of the original participants were excluded due to missing task data, which led to the reanalysis of resting-state fixation data to align participant groupings. Resting-state fixation and task-regressed data were used as inputs in the MS-HBM to estimate precision brain networks for each individual, employing the same standard fcMRI network analysis as outlined in Du et al. (2024). The models were estimated for the resting-state acquisitions and separately for the task acquisitions.

First, the functional connectivity profile of each vertex on the fsaverage6 cortical surface was calculated based on its functional connectivity to 1,175 ROIs uniformly distributed across the fsaverage surface (Yeo et al. 2011). For each run of task-based data, Pearson’s correlation coefficients between the fMRI time series at each vertex (40,962 vertices / hemisphere) and the 1,175 ROIs were computed. The resulting 40,962 × 1,175 correlation matrix was then binarized by keeping the top 10% of the correlations to obtain the functional connectivity profiles (Yeo et al. 2011). Next, the expectation-maximization algorithm for estimating parameters in the MS-HBM was initialized with a 15-network group-level parcellation, ‘DU15NET-Prior’, derived from a subset of the HCP S900 data release (Du et al. 2024) and used to generate within-individual precision network estimates.

After obtaining network estimates separately for resting-fixation and task-based acquisitions, we examined correspondence using qualitative as well as quantitative procedures. Qualitative correspondence was visualized by comparing the network assignments visually on the fully inflated flatmaps so that all similarities and differences could be appreciated. Quantitative correspondence was computed as the percentage of overlap of the assignment of every individual network for each participant. This is a conservative estimate because there are 15 possible network assignments for every vertex. Positive overlap was recorded only when the vertex was assigned to the exact same network between the two compared maps. Moreover, we did not exclude any vertices. Overlap misses, even in regions of low signal-to-noise, penalize the estimates of correspondence. To anchor the interpretation of the estimate, we also compute quantitative similarly for identically acquired resting-state fixation data.

### Model-Free Seed-Region Tests of Spatial Correspondence

Winner-take-all assignments to individual networks do not always capture the complexity of the spatial correlation patterns given vertices assigned to one network may nonetheless have correlations with others. As another means to visualize correspondence between network estimates derived from traditional resting-state fixation versus task acquisitions, a further set of seed-region based analyses was performed.

For each participant, a flatmap was created with the outlines of each network derived solely from the resting-state fixation data. Seed regions were then manually placed in posterior medial, parietal, and prefrontal zones for each network. The correlation patterns from these seed regions from the task-regressed data were then visualized in relation to the network borders. That is, the spatial correlation patterns from the task-regressed data were plotted in relation to the network borders estimated from the traditional resting-state fixation data. This is an extreme test because the task-regressed data are independent and unconstrained by any model assumptions.

### Construct Validity Test of Functional Specificity Using Independent Task Data

To investigate the practical effects of using network estimates derived from task-regressed data, we explored whether the estimated networks could prospectively predict functional response properties in independent tasks collected from separate MRI acquisitions. The logic of this analysis is that network estimates are often used as functional localizers to then predict task responses. For example, the estimate of the LANG network predicts regions preferentially responsive to meaning-based sentence processing tasks (e.g., Braga et al. 2020; Fedorenko, Levy, and Regev 2024). That is, the networks are hypothesized to localize meaningful biological regions that predict response patterns in additional tasks and experimental contexts. For this analysis, we generated network estimates for each participant from task-regressed data that included the following tasks: MOTOR, ODDBALL, NBACK and VISME (referred to as “Task Acquisition” network estimates; see Table 2). These task-based network estimates were directly compared to network estimates derived solely from traditional resting-state fixation data (referred to as “Fixation Acquisition” network estimates). These two sets of estimates -- the Task Acquisition network estimates and the Fixation Acquisition network estimates -- were then used as the basis for testing a triple functional dissociation in independent task data.

There are three well characterized association networks that are juxtaposed with one another across the distributed zones of association cortex that are functionally specialized (DN-A, DN-B, and LANG). These networks are spatially intertwined, making them challenging to separate. Nonetheless, using within-individual precision approaches, they have been reliably identified and functionally dissociated (DiNicola, Braga, and Buckner 2020; Braga et al. 2020; DiNicola, Sun and Buckner 2023; Du et al. 2024). This established and robust functional specificity was leveraged here to test the validity of the network estimates that were derived from the task data. Specifically, we asked whether the Fixation Acquisition network estimates could predict functional responses that are preferential for the EPRJ task contrast (where the response in DN-A is greatest), the ToM task contrasts (where the response in DN-B is greatest), and the SENT task contrast (where the response in LANG is greatest). That is, could network estimates derived exclusively from task data predict a known functional triple dissociation in independent data?

For this final test, functional task data (EPRJ, ToM and SENT) were analyzed using the GLM (FEAT, FSL; Woolrich et al. 2001; Smith et al. 2004) as previously described in Du et al. (2024). GLM outputs included *β*-values for each contrast at each vertex, which were converted to *z*-values within FEAT. For each participant, *z*-value maps from all runs were averaged using *fslmaths* to create a single cross-session map for each task contrast of interest. To quantify the functional response within the independently defined cerebral networks, we calculated the average *z*-value of a specific task contrast for all vertices within each defined network. We then calculated the mean *z*-values across runs to obtain a single response value for each network from each participant. The mean *z*-values across participants were then visualized in the bar graphs, along with the standard error of the mean. In all cases, the networks were defined within individuals based on functional connectivity derived from either the resting-state fixation data (“Fixation Acquisition”) or the separate task-based data (“Task Acquisition”) data. Critically, the target task data used to test for functional dissociation was independent from the data used to estimate the networks.

### Software and Statistical Analysis

Functional connectivity between brain regions was calculated in MATLAB (version 2019a; http://www.mathworks.com; MathWorks, Natick, MA) using Pearson’s product moment correlations. FreeSurfer v6.0.0, FSL, and AFNI were used during data processing. Model-free seed-region analyses were performed using Connectome Workbench v1.3.2. All statistical analyses were performed using R v3.6.2. Network parcellation was performed using code from Kong et al. (2019) on GitHub (https://github.com/ThomasYeoLab/CBIG/tree/master/stable_projects/brain_parcellation/Kong2019_MSHBM).

## Data Availability

Individual participant data are available in the NIH repository (https://nda.nih.gov). Parcellation, SNR and task contrast maps related to this manuscript are available on Balsa (https://balsa.wustl.edu/study/zK166).

Task descriptions, contrast descriptions, and code are provided on Harvard Dataverse (https://doi.org/10.7910/DVN/AVB4BW).

## Acknowledgments

We thank the Harvard Center for Brain Science neuroimaging core and FAS Division of Research Computing. We thank T. O’Keefe for assisting in optimization of data processing, R. Mair for MRI physics support, and N. Saadon-Grosman, H. Kosakowski, W. Sun, V. Tripathi, A. Billot, L. DiNicola, and L. Hanford for discussion. The multi-band EPI sequence was generously provided by the Center for Magnetic Resonance Research (CMRR) at the University of Minnesota.

